## SupplementalFigures for "Hsp70 is phosphorylated in a conserved response to DNA damage and contributes to cell cycle control"

### SUPPLEMENTAL FIGURES

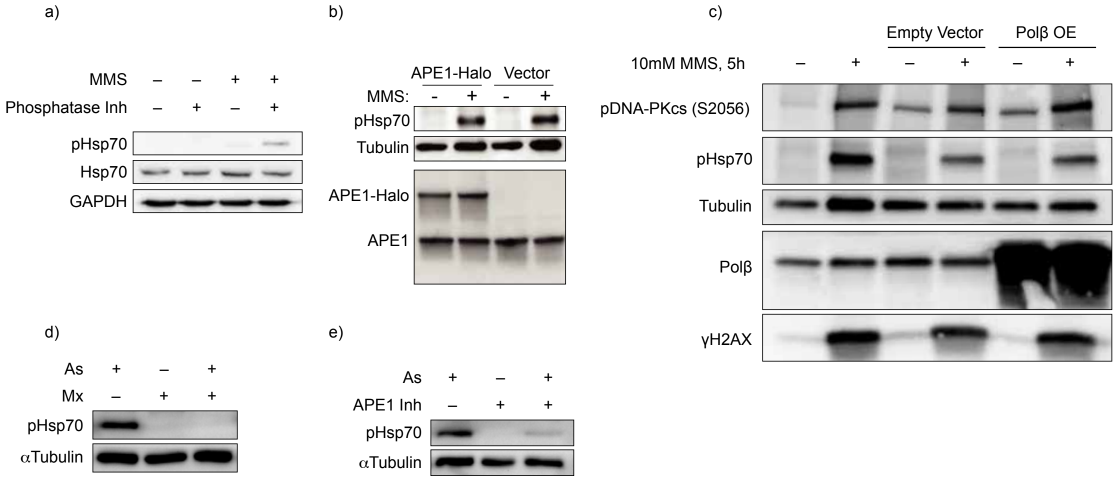


**Figure S1 (related to Figure 2). Base excision repair drives phosphorylation of Hsp70 in human cells**

**a)** MMS-induced pHsp70 band is sensitive to phosphatases. Cells were treated with 10 mM MMS for 5 h or left untreated, and then lysed in the presence or absence of phosphatase inhibitor. pHsp70, Hsp70, and GAPDH levels were examined via immunoblotting. Data are representative of n = 3 independent experiments. **b)** Effect of overexpression of APE1 on pHsp70 levels. Cells were transiently transfected with APE1-Halo or EV overnight before treatment with MMS. pHsp70 and APE1 levels were examined via immunoblotting, with tubulin as a loading control. Data are representative of n = 3 independent experiments. **c)** Effect of overexpression of Polβ on pHsp70 levels. Cells were transiently transfected with Polβ or EV overnight before treatment with MMS. pHsp70 and Polβ levels were examined via immunoblot. γH2AX was probed to detect overall levels of DNA damage. DNA-PKcs autoactivation were monitored by immunoblotting pDNA-PKcs (S2056). Tubulin was used as a loading control. Data are representative of n = 2 independent experiments. **d)** Masking of AP sites prevents arsenite-induced pHsp70 accumulation. Cells were pretreated with 60 mM methoxyamine (Mx) for 30 min, followed by cotreatment with 30 mM Mx and 0.5 mM sodium arsenite for 5 h. pHsp70 and the loading control α tubulin were detected by immunoblotting. Data are from n = 1 experiment. **e)** Inhibition of APE1 reduces arsenite-induced pHsc70. Cells were pretreated for 1 h with 50 μM APE1 inhibitor (APE1 compound III) before arsenite treatment. pHsp70 and the loading control α–tubulin were detected by immunoblotting. Data are from n = 1 experiment.

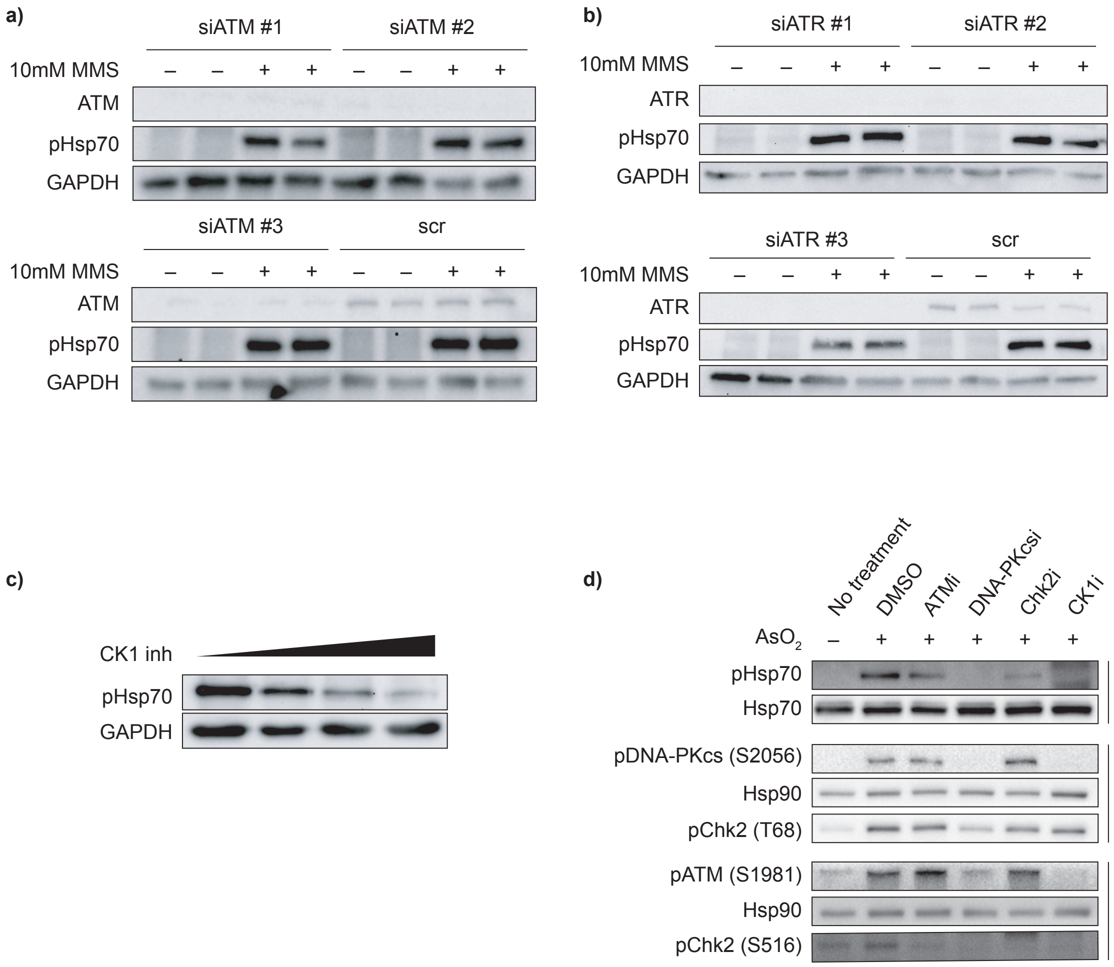


**Figure S2 (related to Figure 3). DDR kinase activity is upstream of Hsp70 phosphorylation**
**a)** ATM knockdown does not reduce pHsp70 levels during MMS treatment. Cells were transiently transfected with three independent siRNAs targeting ATM or a scramble control for 72 h, followed by treatment with 10 mM MMS for 5 h. ATM, pHsp70, and GAPDH (loading control) were detected by immunoblotting. Data are from n = 1 experiment. **b)** ATR knockdown does not reduce pHsp70 levels during MMS treatment. Cells were transiently transfected with three independent siRNAs targeting ATR or a scramble control for 72 h, followed by treatment with 10 mM MMS for 5 h. ATR, pHsp70, and GAPDH (loading control) were detected by immunoblotting. Data are from n = 1 experiment. **c)** CK1 inhibition leads to dose dependent decrease of pHsp70 accumulation. Cells were pretreated with vehicle or the CK1 inhibitor PF-670462 (5 μM, 25 μM, or 50 μM) for 1 h, followed by 10 mM MMS treatment for 5 h. pHsp70 and the loading control GAPDH were detected by immunoblotting. Data are representative of n = 3 independent experiments. Data are representative of n = 3 independent experiments. **d)** Pharmacological inhibition of ATM, DNA-PKcs, Chk2, and CK1 decrease Hsp70 phosphorylation during arsenite treatment. Cells were pretreated for 1h with inhibitors for ATM (200 nM AZD1390), DNA-PKcs (2 μM AZD7648), Chk2 (5 μM CCT241533), CK1(50 μM PF-670462) or with vehicle control (DMSO), then treated with 0.5 mM arsenite for 5h. Immunoblotting was performed against pHsp70, active DDR kinases (pDNA-PKcs(S2056), pChk2(T68), pATM(S1981), pChk2(S516)), or the loading controls Hsp70 or Hsp90. Data are representative of n = 2 independent experiments.

a)

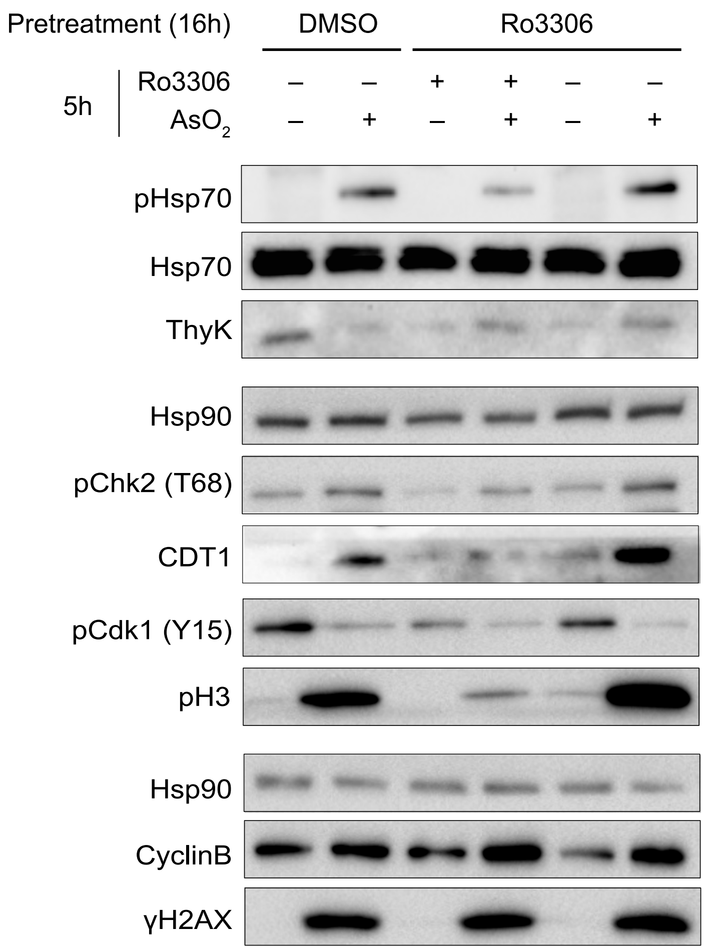


**Figure S3 (related to Figure 4). Mitosis precedes Hsp70 phosphorylation**

**a)** G2/M stalling by CDK1 inhibition reduces arsenite induced pHsp70 levels. Cells were pretreated with 10 μM CDK1 inhibitor Ro3306 or DMSO for 16 h, washed twice with PBS, then treated again with Ro3306 or DMSO in the presence or absence of 0.5 mM sodium arsenite for 5 h. Immunoblotting was performed for pHsp70, cell cycle markers (pCdk1 Y15, pH3, CDT1, cyclin B, ThyK), DDR markers (pChk2 T68, γH2AX), and loading controls Hsp70 and Hsp90. Data are representative of n = 2 independent experiments.

a)

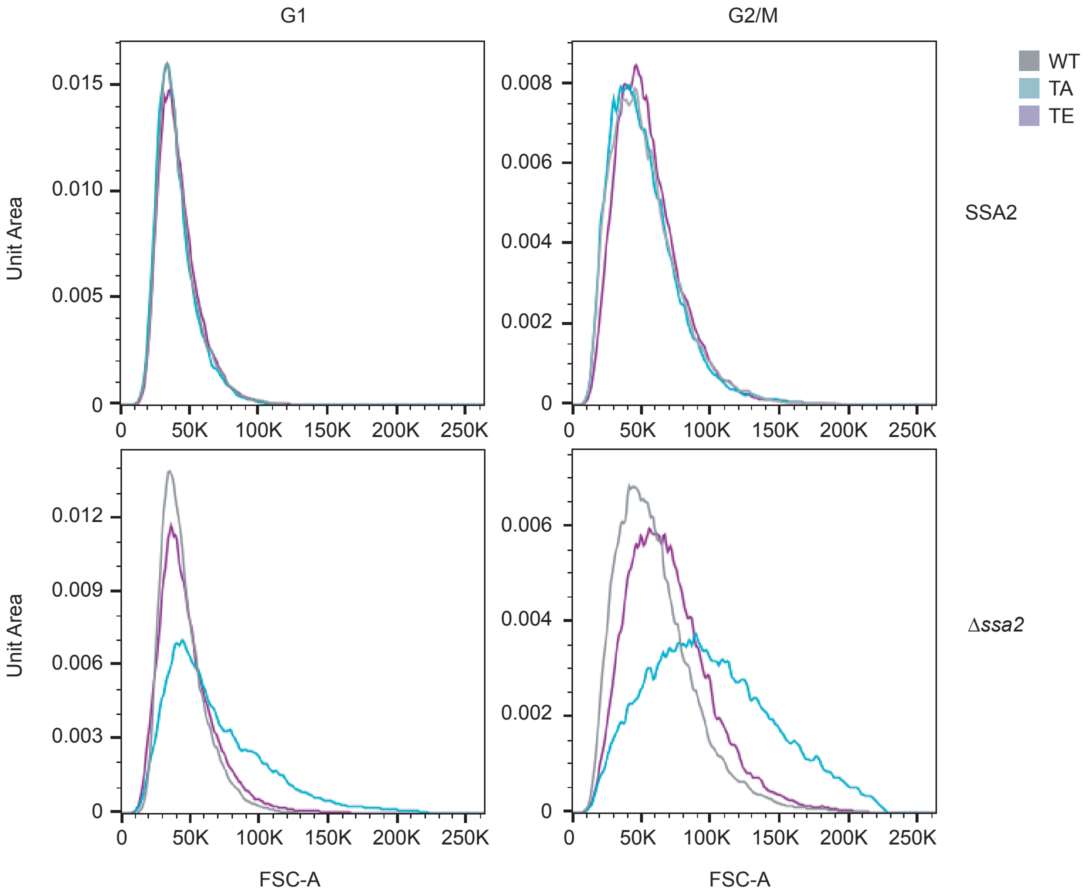


**Figure S4 (related to Figure 5). Cell size is increased for Ssa1 phosphomutant yeast strains in the absence of Ssa2**

**a)** Size distribution of Ssa1 mutant strains. Yeast were grown to mid-log phase, adjusted to the same concentration, and immediately fixed. Cells were stained with Sytox Green and analyzed by flow cytometry to determine DNA content. Cells were gated by singlets and then by DNA content (1N = G1; 2N = G2/M). Size is displayed as forward scatter area (FSC-A) on the X axis and count normalized to unit area on the Y axis. The left column are strains in the SSA2 background, and the right is strains in the *ssa2*Δ background. Ssa1 mutants are color coded (WT gray, TA blue, TE purple). Data are representative of n = 4 independent technical replicates with 2 biological replicates per strain.
